# Infection and transmission of SARS-CoV-2 depends on heparan sulfate proteoglycans

**DOI:** 10.1101/2020.08.18.255810

**Authors:** Marta Bermejo-Jambrina, Julia Eder, Tanja M. Kaptein, John L. van Hamme, Leanne C. Helgers, Killian E. Vlaming, Philip J.M. Brouwer, Alexander P.J. Vlaar, Frank E.H.P. van Baarle, Marcel Spaargaren, Godelieve J. de Bree, Bernadien M. Nijmeijer, Neeltje A. Kootstra, Marit J. van Gils, Rogier W. Sanders, Teunis B. H. Geijtenbeek

**Author notes:** Corresponding author: (TBHG). These authors contributed equally (Marta Bermejo-Jambrina, Julia Eder).

## Abstract

The current pandemic caused by severe acute respiratory syndrome coronavirus-2 (SARS-CoV-2) and outbreaks of new variants highlight the need for preventive treatments. Here we identified heparan sulfate proteoglycans as attachment receptors for SARS-CoV-2. Notably, neutralizing antibodies against SARS-CoV-2 isolated from COVID-19 patients interfered with SARS-CoV-2 binding to heparan sulfate proteoglycans, which might be an additional mechanism of antibodies to neutralize infection. SARS-CoV-2 binding to and infection of epithelial cells was blocked by low molecular weight heparins (LMWH). Although dendritic cells (DCs) and mucosal Langerhans cells (LCs) were not infected by SARS-CoV-2, both DC subsets efficiently captured SARS-CoV-2 via heparan sulfate proteoglycans, and transmitted the virus to ACE2-positive cells. Moreover, human primary nasal cells were infected by SARS-CoV-2 and infection was blocked by pre-treatment with LMWH. These data strongly suggest that heparan sulfate proteoglycans are important attachment receptors facilitating infection and transmission, and support the use of LMWH as prophylaxis against SARS-CoV-2 infection.

## Introduction

Severe acute respiratory syndrome coronavirus 2 (SARS-CoV-2) emerged in Wuhan, China in late 2019 and can cause coronavirus disease 2019 (COVID-19), an influenza-like disease ranging from mild respiratory symptoms to severe lung injury, multi organ failure and death (Yuki *et al*, 2020; Zhou *et al*, 2020; Zhu *et al*, 2020). SARS-CoV-2 spread quickly and has caused a pandemic with a severe impact on global health and world economy (Nicola *et al*, 2020; World Health Organization, 2020b). SARS-CoV-2 is transmitted predominantly via large droplets expelled from the upper respiratory tract through sneezing and coughing (Ferioli *et al*, 2020; Harapan *et al*, 2020) and is subsequently taken up via mucosal surfaces of the nose, mouth and eyes (Peiris *et al*, 2003). SARS-CoV-2 infects epithelial cells in the respiratory tract, such as ciliated mucus secreting bronchial epithelial cells and type 1 pneumocytes in the lung, as well as epithelial cells in the gastrointestinal tract (Hui *et al*, 2020; Lamers *et al*, 2020). For more than a year lockdown strategies and social distancing have been used to mitigate viral spread but due to negative socioeconomic consequences these are not feasible long-term solutions (Brooks *et al*, 2020; Wright *et al*, 2020). Currently, several COVID-19 vaccines have been developed and worldwide vaccination programs have been initiated (Mathieu *et al*, 2021), which aim to curb and stop the pandemic. However, immunocompromised individuals as well as people on immunosuppressive drugs are potentially less protected by vaccinations (Agha *et al*, 2021; Boyarsky *et al*, 2021). Moreover, current vaccine candidates might be less effective against new SARS-CoV-2 variants (Collier *et al*, 2021; Wang *et al*, 2021). Thus, there is a need for protective strategies specifically targeting SARS-CoV-2 to prevent further dissemination.

SARS-CoV-2 belongs to the betacoronaviruses, a family that also includes SARS-CoV and MERS-CoV (Letko *et al*, 2020). The coronavirus Spike (S) protein is a class I fusion protein that mediates virus entry (Bosch *et al*, 2003; Hulswit *et al*, 2016). The S protein consist of two subunits; S1 directly engages via its receptor-binding domain (RBD) with host surface receptors (Li *et al*, 2005; Wang *et al*, 2013) and S2 mediates fusion between virus and cell membrane (Burkard *et al*, 2014; Xia *et al*, 2020). SARS-CoV-2 uses angiotensin-converting enzyme 2 (ACE2) as its main receptor (Hoffmann *et al*, 2020; Letko *et al*., 2020). ACE2 is a type I integral membrane protein abundantly expressed on epithelial cells lining the respiratory tract (Hamming *et al*, 2004) but also the ileum, esophagus and liver (Zou *et al*, 2020b) and ACE2 expression dictates SARS-CoV-2 tropism (Lamers *et al*., 2020). However, it remains unclear whether SARS-CoV2 requires other receptors for virus entry. Neutralizing monoclonal antibodies against SARS-CoV-2 have been identified that are directed not only at the RBD but also outside the RBD (Brouwer *et al*, 2020), suggesting that other mechanisms of neutralization or other (co-)receptors might be involved.

Heparan sulfates are expressed by most cells including epithelial cells as heparan sulfate proteoglycans and these have been shown to interact with viruses such as HIV-1, HCV, Sindbis virus and also SARS-CoV (Byrnes & Griffin, 1998; Jiang *et al*, 2012; Milewska *et al*, 2014; Nijmeijer *et al*, 2020; Roderiquez *et al*, 1995). Recently, it was shown that the S protein of SARS-CoV-2 interacts with heparan sulfates, which might be required for infection (Clausen *et al*, 2020; Zhang *et al*, 2020).

Here we show that the heparan sulfate proteoglycans are important for infection of polarized epithelial cells as well as primary nasal cells with SARS-CoV-2. Infection is inhibited by heparin and low molecular weight heparins (LMWH). Mucosal dendritic cell subsets captured SARS-CoV-2 via heparan sulfate proteoglycans. The different DC subsets did not become infected but transmitted SARS-CoV-2 to ACE2-positive cells, which might facilitate virus dissemination. Our findings suggest that heparan sulfate proteoglycans function as attachment receptors for SARS-CoV-2 and LMWH can be used as prophylactics against SARS-CoV-2 or prevent dissemination early after infection.

## Results

### SARS-CoV-2 binds to heparan sulfates expressed by cells

We incubated Huh 7.5 cells that express ACE2 (Fig EV1A) with SARS-CoV-2 pseudovirus, which consists of HIV-1 particles pseudotyped with SARS-CoV-2 S protein (Brouwer *et al*., 2020). Virus binding was determined by measuring HIV-1 p24 uptake by ELISA. SARS-CoV-2 pseudovirus attached to Huh 7.5 cells, which was blocked by anti-ACE2 antibodies (Brouwer *et al*., 2020) as well as by neutralizing antibodies from COVID-19 patients (Brouwer *et al*., 2020) (Fig 1A). Unfractionated (UF) heparin inhibited binding of SARS-CoV-2 pseudovirus to Huh 7.5 cells comparable to the neutralizing or anti-ACE2 antibodies (Fig 1A). Enzymatic removal of heparan sulfates on the cell surface by Heparinase treatment decreased SARS-CoV-2 virus binding (Fig 1B and Fig EV1A). Exostosin-1 (EXT1) knockdown strongly decreased expression of heparan sulfates on the cell surface (Ren *et al*, 2018) (Fig 1C). SARS-CoV-2 pseudovirus attached to XG1 cells, which was blocked by UF Heparin, whereas knockdown of EXT1 abrogated SARS-CoV-2 pseudovirus binding (Fig 1D). These data suggest that heparan sulfates are important for attachment of SARS-CoV-2 to cells.

**Fig 1.**
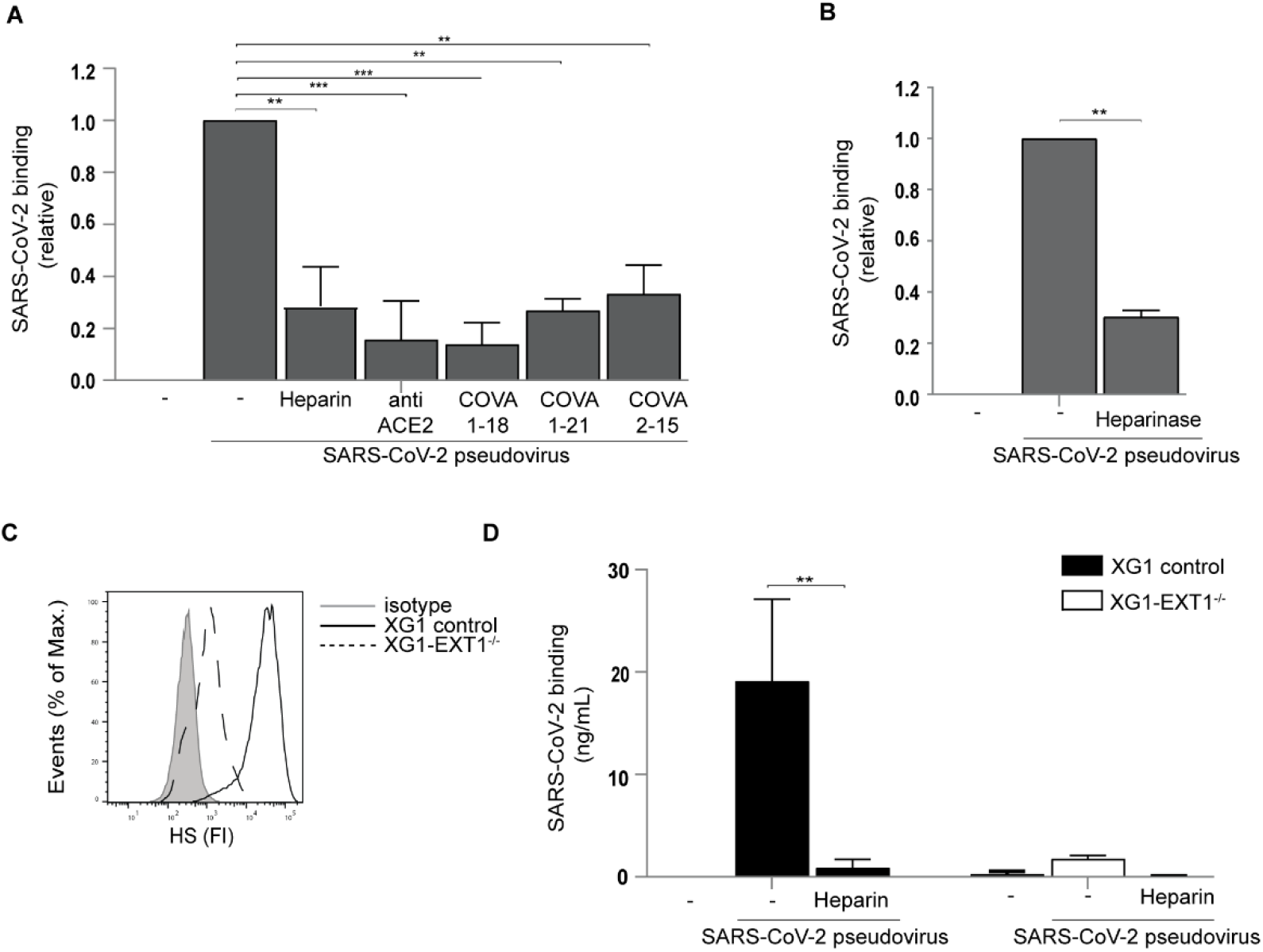
SARS-CoV-2 binds to heparan sulfates expressed by cells. (A) Huh 7.5 cells were preincubated with neutralizing antibody to ACE2 and SARS-CoV-2 pseudovirus was pre-incubated with mAb COVA1-18, COVA1-21 and COVA2-15 or UF heparin (250IU). Cells were incubated with SARS-CoV-2 pseudovirus and binding was determined by ELISA. (B) Heparan sulfates were removed from Huh 7.5 cells by heparinase treatment. SARS-CoV-2 pseudovirus binding was determined by ELISA. (C) Flow cytometry analysis of cell surface expression of heparan sulfates (HS) in control transduced cells or upon CRISPR/Cas9-mediated EXT1 KO (EXT1-/-). (D) XG1 cells (control and EXT1-/-) were exposed to SARS-CoV-2 pseudovirus and binding was measured by ELISA. Data show the mean values and error bars are the SEM. Statistical analysis was performed using (A) ordinary one-way ANOVA with Tukey multiple-comparison test. **p= 0.0010, ***p= 0.0006, ***p= 0.0005, **p= 0.0018, **p= 0.0036 (n = 3), (B) two-tailed, unpaired Student’s t-test with Welch’s correction. **p= 0.0012 (n = 3), (D) two-way ANOVA with Dunnett’s multiple-comparison test. **p= 0.0013 (n=3). RLU: relative light units.

### Low molecular weight heparins inhibit SARS-CoV-2 infection

To determine the effect of UF Heparin on SARS-CoV-2 infection, we infected Huh 7.5 cells with SARS-CoV-2 pseudovirus, expressing the luciferase reporter gene, and determined infection by luciferase activity. UF Heparin blocked infection in a dose-dependent manner (Fig 2A). Low molecular weight heparin (LMWH) have replaced UF heparin in the clinic as anti-coagulant treatment due to their smaller size and superior pharmacological properties (Kakkar, 2004). LMWH enoxaparin blocked SARS-CoV-2 pseudovirus infection in a dose dependent manner to similar levels as UF Heparin (Fig 2A) without affecting cell viability of Huh 7.5 cells (Fig EV1E). Not only enoxaparin but also other clinically approved LMWH blocked binding of SARS-CoV-2 pseudovirus to Huh 7.5 cells (Fig 2B). The different LMWH also blocked infection of Huh 7.5 cells with SARS-CoV-2 pseudovirus to a similar extent as enoxaparin (Fig 2C).

**Fig 2.**
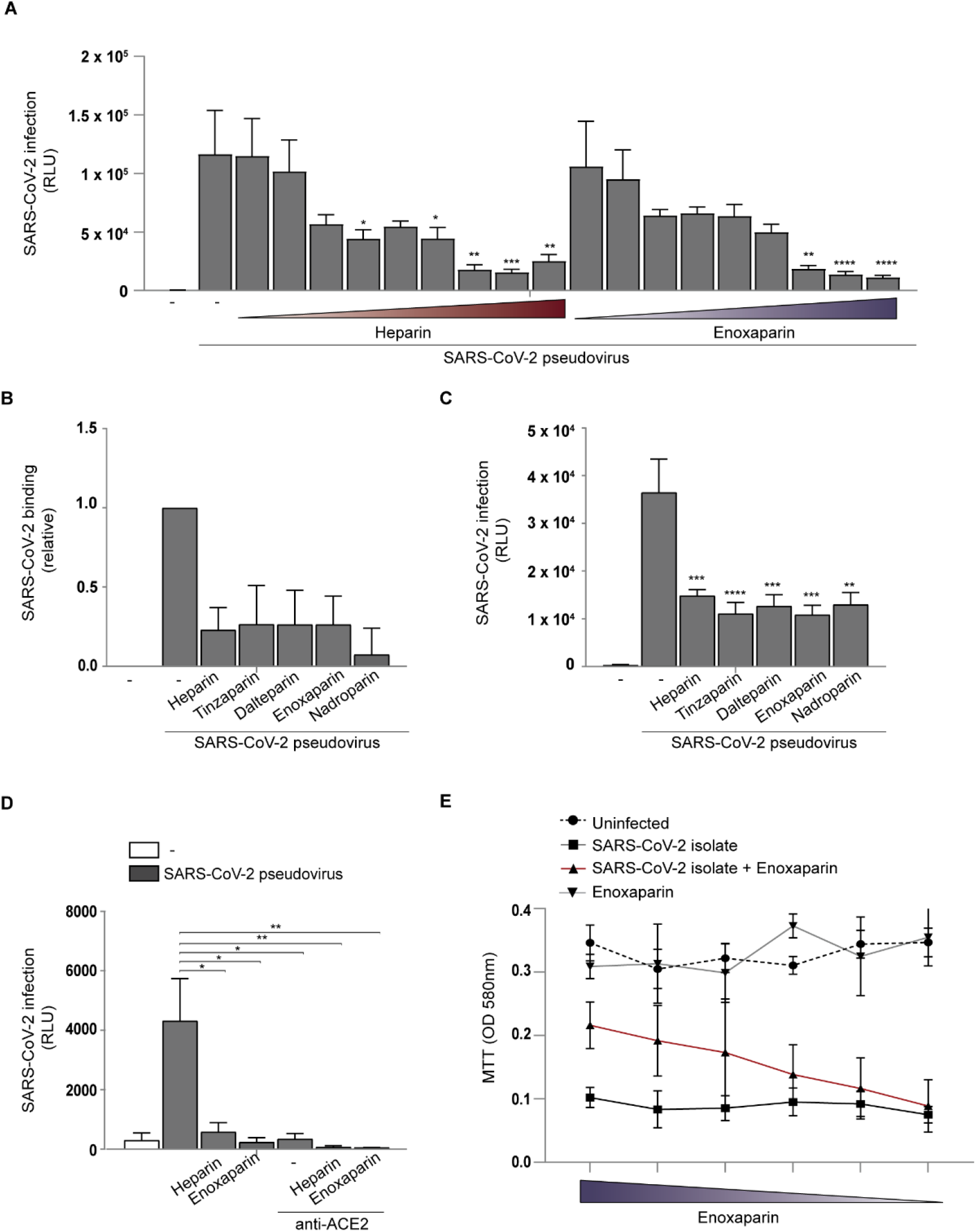
Low molecular weight heparins inhibit SARS-CoV-2 infection. (A) Huh7.5 cells were exposed with SARS-CoV-2 pseudovirus pretreated with different concentrations (0.0001IU-250IU) of UF heparin and enoxaparin. Infection was determined by luciferase assay. (B) SARS-CoV-2 pseudovirus binding to Huh7.5 after 4 hours was determined by ELISA. (C) SARS-CoV-2 pseudovirus infection of Huh7.5 cells was measured after 5 days by luciferase assay. (D) 293T cells expressing ACE2 were infected with SARS-CoV-2 in presence or absence of antibodies against ACE2, UF heparin (250IU) or enoxaparin (250IU). (E) VeroE6 cells were infected with a SARS-CoV-2 isolate (Italy TCID50 104) previously treated with serial dilutions of enoxaparin. MTT assay detects live cells, and the loss of MTT staining as determined by spectrometer (OD 580nm) is indicative of SARS-CoV-2 cytotoxicity. Data show the mean values and error bars are the SEM. Statistical analysis was performed using (A) ordinary one-way ANOVA with Dunnett’s multiple-comparison test. *p= 0.0044, **p= 0.0012, ***p= 0.0003, **p= 0.0037, **p= 0.0014, ****p= 0.0007, ****p= 0.0007. (B) ordinary one-way ANOVA with Tukey’s multiple-comparison test. (n = 3), (C) ordinary one-way ANOVA with Dunnett’s multiple-comparison test. **p= 0.0018, ***p= 0.0003, ***p= 0.0004, ****p< 0.0001, **p= 0.0036, **p= 0.0012, **p= 0.0029 (n = 3 donors measured in triplicate), (D) ordinary one-way ANOVA with Dunnett’s multiple-comparison test. *p= 0.0380, *p= 0.0136, *p= 0.0189, **p= 0.0080, **p= 0.0074 (n = 3 measured in triplicates). RLU: relative light units.

Next we investigated whether ACE2 is required for infection in presence of heparan sulfates. Human kidney epithelial 293T cells were not susceptible to SARS-CoV-2 pseudovirus whereas ectopic expression of ACE2 rendered these cells susceptible to SARS-CoV-2 pseudovirus (Fig 2D and Fig EV1B and D). Infection was abrogated by both LMWH enoxaparin and UF heparin to a similar level as antibodies against ACE2 (Fig 2D). The combination of ACE2 antibodies and LMWH enoxaparin or UF heparin blocked infection of 293T-ACE2 cells (Fig 2D). These data suggest that heparan sulfates act as attachment receptors that allow the virus to bind to cells for infection via ACE-2.

Simian Vero E6 cells are highly susceptible to SARS-CoV-2, which causes severe cytopathic effects (CPE) (Zhou *et al*., 2020). We used a tetrazolium dye (MTT) colorimetric cell viability assay (Präbst *et al*, 2017) to investigate the role of LMWH upon SARS-CoV-2 infection. Within 48 hours of VeroE6 cell inoculation with a primary SARS-CoV-2 isolate (hCoV-19/Italy) severe CPE was observed and cell viability strongly decreased (Fig 2E), which was counter-acted by LMWH enoxaparin in a concentration dependent manner. These data support an important role for heparan sulfates to facilitate SARS-CoV-2 infection via ACE-2.

### SARS-CoV-2 infection of polarized epithelial cells is blocked by UF heparin and LMWH

The colorectal adenocarcinoma Caco-2 and bronchial adenocarcinoma Calu-3 cells represent models for human intestinal and respiratory epithelial cells, respectively (Artursson *et al*, 2001; Harcourt *et al*, 2011). Both cell lines were cultured on microporous filters with an air liquid interface to achieve a polarized monolayer and polarization was monitored by transepithelial electrical resistance (TEER). TEER increased over time to confirm that the cells are polarized after culture of more than 14 days (Fig 3A). Undifferentiated Caco-2 and Calu-3 expressed low levels of ACE2 but polarization of the cells by strongly increased ACE2 expression (Fig 3B). Polarized Caco-2 cells as well as Calu-3 cells strongly bound to SARS-CoV-2 pseudovirus, which was significantly blocked by LMWH to similar levels as antibodies against ACE2 (Fig 3C and D). The combination of an antibody against ACE2 and enoxaparin did not further decrease binding. Next we exposed the polarized Calu-3 cells to the primary SARS-CoV-2 isolate for 24 hours and after extensive washing cells were cultured for another 24 hours. Infection was measured by qPCR of both supernatant and cell-lysates for SARS-CoV-2 ORF1b transcripts to determined infection. Calu-3 polarized cells were productively infected by the SARS-CoV-2 isolate as shown by the SARS-CoV-2 ORF1b transcripts in both cell-lysate and supernatant (Fig 3E and F). Notably, productive infection of Calu-3 cells was inhibited by LMWH to a similar level as ACE2 antibodies (Fig 3E and F). These data suggest that heparan sulfates are required for binding and infection of polarized respiratory epithelial cells with SARS-CoV-2 pseudovirus as well as primary SARS-CoV-2 isolate.

**Fig 3.**
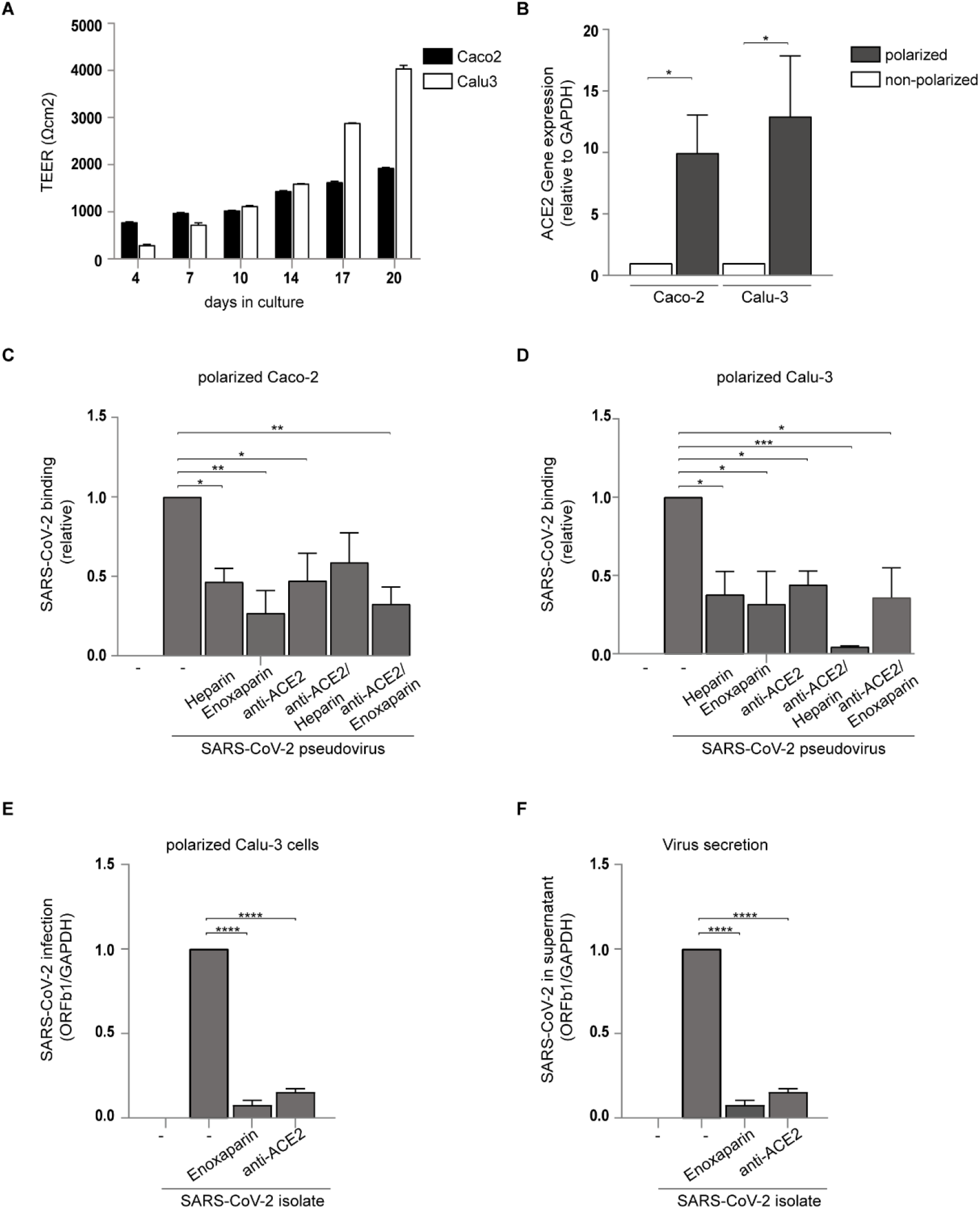
SARS-CoV-2 infection of polarized epithelial cells is blocked by UF heparin and LMWH. (A) Barrier integrity was analysed by TEER measurements of Caco-2 and Calu-3 over a period of 20 days. (B) ACE2 cell surface expression on Caco-2 and Calu-3 was determined by quantitative real-time PCR. One representative donor out of 3 is depicted. (C-D) SARS-CoV-2 binding was measured in polarized Caco-2 (C) and Calu-3 (D) cells in presence of antibodies against ACE2, and UF heparin (250IU) or enoxaparin (250IU). (E) Polarized Calu-3 were infected with a SARS-CoV-2 isolate (Italy TCID50 104) in presence of antibodies against ACE2 and enoxaparin (250IU). Virus was detected after lysing by quantitative real-time PCR of viral RNA (E) and virus production was determined by detecting viral RNA in supernatant using quantitative real-time PCR (F). Data show the mean values and error bars are the SEM. Statistical analysis was performed using (B) ordinary one-way ANOVA with Tukey’s multiple-comparison test. ****p < 0.0001 (n = 1), (C-D) ordinary one-way ANOVA with Tukey’s multiple-comparison test. *p= 0.00466, **p= 0.0069, *p= 0.0181, **p= 0.0046 (n=3 Caco-2 donors measured in triplicate), *p= 0.0262, *p= 0.0149, *p= 0.0475, ***p= 0.0006, *p= 0.0221 (n=3 Calu-3 donors measured in triplicate), (E-F) ordinary one-way ANOVA with Tukey’s multiple-comparison test. ****p<0.0001. TEER: Transepithelial electrical resistance, RLU: relative light units.

### Heparan sulfate proteoglycans Syndecan 1 and 4 are important for SARS-CoV-2 binding

The heparan sulfate proteoglycan family of Syndecans are particularly important in facilitating cell adhesion of several viruses (Bacsa *et al*, 2011; Nijmeijer *et al*., 2020). Therefore, we used the Namalwa B cell-line that ectopically expressed Syndecan 1 and 4 (Fig EV2A and EV2B) as these Syndecans are expressed by epithelial cells (Hayashida *et al*, 2006; Teng *et al*, 2012). Namalwa cells did not express ACE2 (Fig EV2C). In contrast to the parental control Namalwa cells, Syndecan 1 and Syndecan 4 expressing cells interacted strongly with SARS-CoV-2 pseudovirus (Fig 4A). Both UF heparin and enoxaparin blocked the interaction of Syndecan 1- and -4 expressing cells with SARS-CoV-2 pseudovirus (Fig 4A). Moreover, the primary SARS-CoV-2 isolate attached to both Syndecan 1 and Syndecan 4 expressing cells, and UF Heparin as well as enoxaparin blocked binding to levels observed for the parental control cells (Fig 4B). These data indicate that Syndecan 1 and 4 are important heparan sulfate proteoglycans involved in SARS-CoV-2 binding and infection.

**Fig 4.**
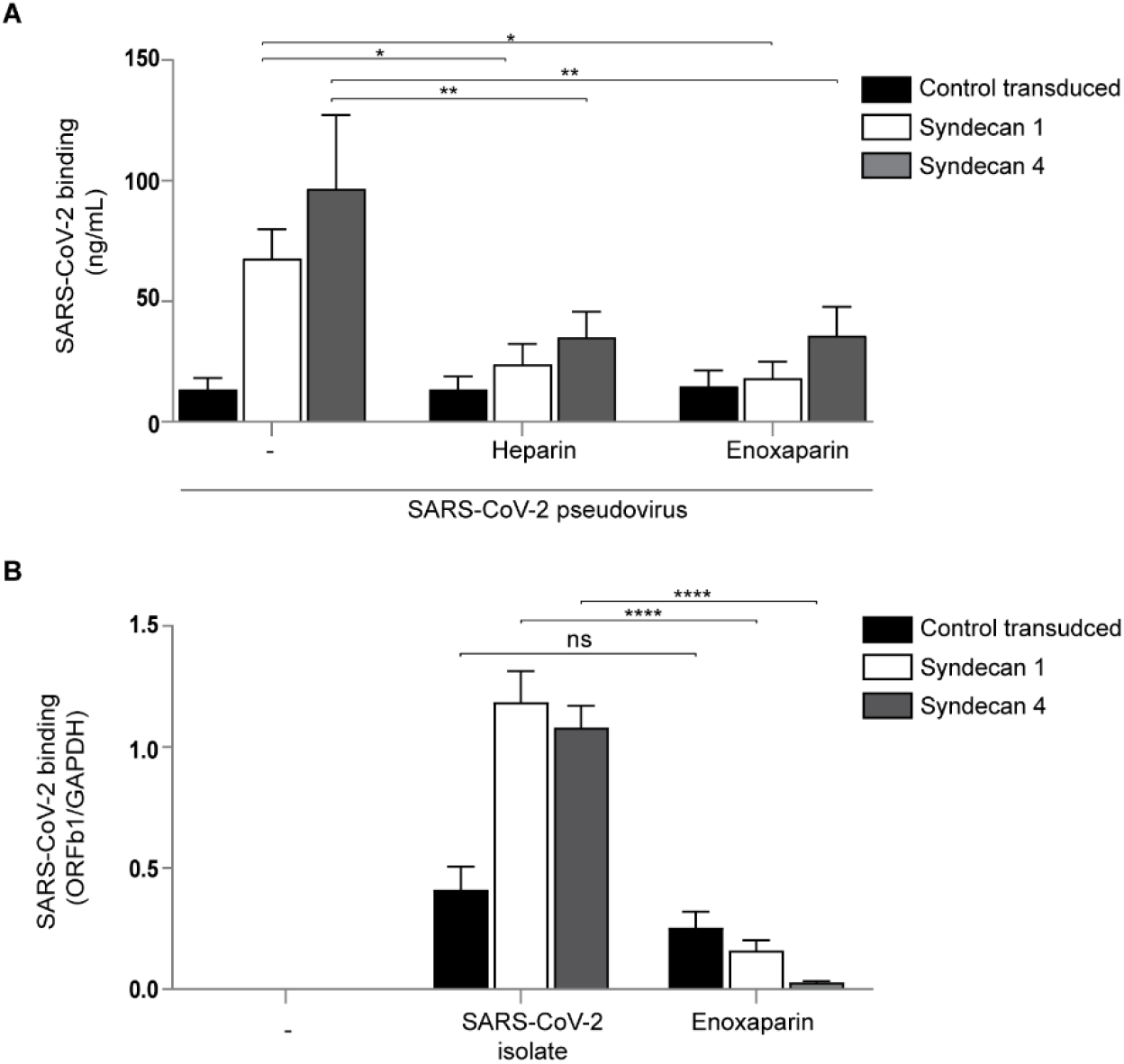
Heparan sulfate proteoglycans Syndecan 1 and 4 are important for SARS-CoV-2 binding. (A-B) Namalwa cells ectopically expressing either Syndecan 1 or 4 were exposed to SARS-CoV-2 pseudovirus (A) or SARS-CoV-2 isolate (Italy TCID50 104) (B) in presence or absence of UF heparin (250IU) or enoxaparin (250IU) and binding was measured after 4h by ELISA or quantitative real-time PCR, respectively. Data show the mean values and error bars are the SEM. Statistical analysis was performed using (A) 2way-ANOVA with Dunnett’s multiple-comparison test. **p= 0.0032, **p= 0.0035, *p= 0.0408, *p= 0.0189 (n = 7), (B) 2way-ANOVA with Sidak’s multiple-comparison test. ****p<0.0001 (n = 3 measured in triplicate). RLU: relative light units.

### Neutralizing antibodies against SARS-CoV-2 interfere with SARS-CoV-2 binding to Syndecan 1

Several antibodies against SARS-CoV-2 were isolated from COVID-19 patients and some of these were potent neutralizing antibodies against SARS-CoV-2 that target the RBD (COVA1-15, COVA1-18) as well as the non-RBD (COVA1-21) of the S protein (Brouwer *et al*., 2020). Therefore, we investigated whether antibodies against SARS-CoV-2 interfere with the interaction of heparan sulfates with SARS-CoV-2. We treated SARS-CoV-2 pseudovirus with different antibodies and measured virus binding to ACE2-negative Syndecan 1-positive Namalwa cells. Notably, only the three neutralizing antibodies against SARS-CoV-2, COVA-1-15, 1-18 and 1-21 blocked the interaction of SARS-CoV-2 pseudovirus with Syndecan 1 in a concentration dependent manner and to similar levels as observed for LMWH (Fig 5A). In contrast, non-neutralizing antibodies did not inhibit virus binding (Fig 5A).

**Fig 5.**
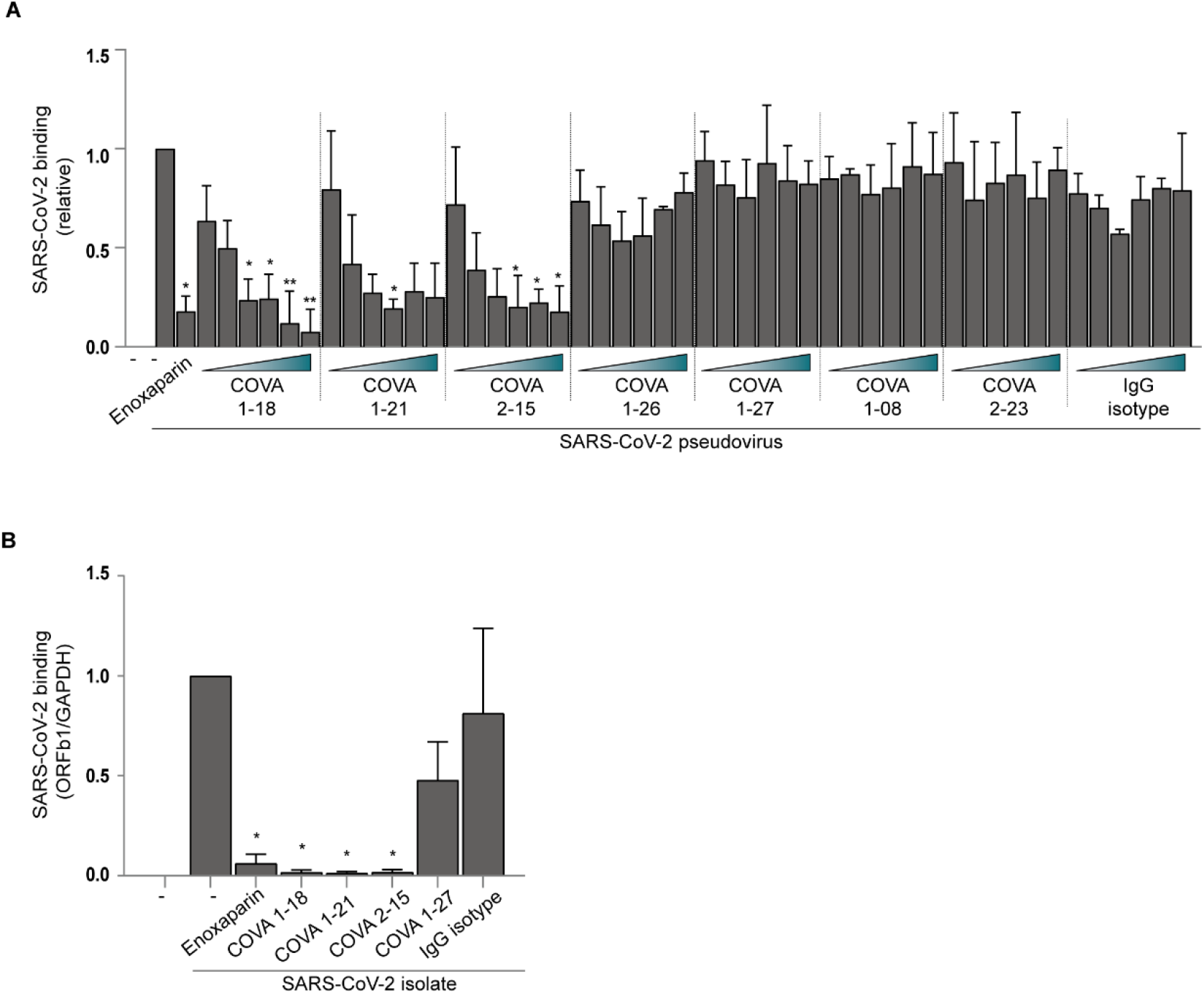
Neutralizing antibodies against SARS-CoV-2 interfere with SARS-CoV-2 binding to Syndecan 1. (A) SARS-CoV-2 pseudovirus was treated with neutralizing antibodies against SARS-CoV-2 (COVA1-18, COVA1-21 and COVA2-15) and isotype control at concentrations 0.0001, 0.0005, 0.001, 0.005, 0.01 and 0.05 µg/mL. Binding to Syndecan 1 expressing cells was determined by ELISA. (B) SARS-CoV-2 isolate was pretreated with LMWH enoxaparin, neutralizing antibodies (COVA 1-18, 1-21 and 2-15) and an isotype control. Detection of viral binding to Syndecan 1 expressing Namalwa was measured by quantitative real-time PCR. Data show the mean values and error bars are the SEM. Statistical analysis was performed using (A) ordinary one-way with Dunnett’s multiple-comparison test. *p= 0.0213, *p= 0.0423, *p= 0.0467, **p= 0.0095, **p= 0.0051, *p= 0.0255, *p= 0.0281, *p= 0.0370, *p= 0.0206 (n = 3), (B) ordinary one-way with Tukey’s multiple-comparison test. *p= 0.0462, *p= 0.0365, *p= 0.0357, *p= 0.0367. (n = 2 measured in duplicates).

Next we determined the ability of the antibodies to block binding of the primary SARS-CoV-2 isolate to Syndecan 1 expressing Namalwa. Similar as observed for SARS-CoV-2 pseudovirus, the three neutralizing COVA antibodies blocked the interaction of the SARS-CoV-2 isolate with Syndecan 1, whereas non-neutralizing antibody COVA-1-27 did not block binding (Fig 5B). These data strongly suggest that neutralizing RBD and non-RBD targeting antibodies against SARS-CoV-2 interfere with SARS-CoV-2 binding to heparan sulfate proteoglycans.

### SARS-CoV-2 targets dendritic cells for dissemination

SARS-CoV-2 infects cells in nasal mucosa, lung and the intestinal tract but mechanisms for dissemination of the virus from the respiratory and intestinal tract are unclear. Different DC subsets are involved in dissemination of various viruses including HIV-1 and SARS-CoV (Jones *et al*, 2008; Marzi *et al*, 2004; Ribeiro *et al*, 2016). We differentiated monocytes to DCs, which is a model for submucosal DC, and also isolated primary human Langerhans cells (LCs) from skin (de Witte *et al*, 2007b; Sarrami-Forooshani *et al*, 2014) as this DC subset resides in squamous mucosa of different tissues including nasal and intestinal mucosa (Merad *et al*, 2008; Nijmeijer *et al*, 2019). Both monocyte-derived DCs and primary LCs bound SARS-CoV-2 pseudovirus and binding was inhibited by UF heparin as well as LMWH enoxaparin (Fig 6A and B). Neither DCs nor LCs were infected by SARS-CoV-2 pseudovirus (Fig 6C), which is due to the absence of ACE2 expression on both subsets (Fig 6D). These data suggest that primary DC subsets capture SARS-CoV-2 via heparan sulfate proteoglycans but this does not lead to infection.

**Fig 6.**
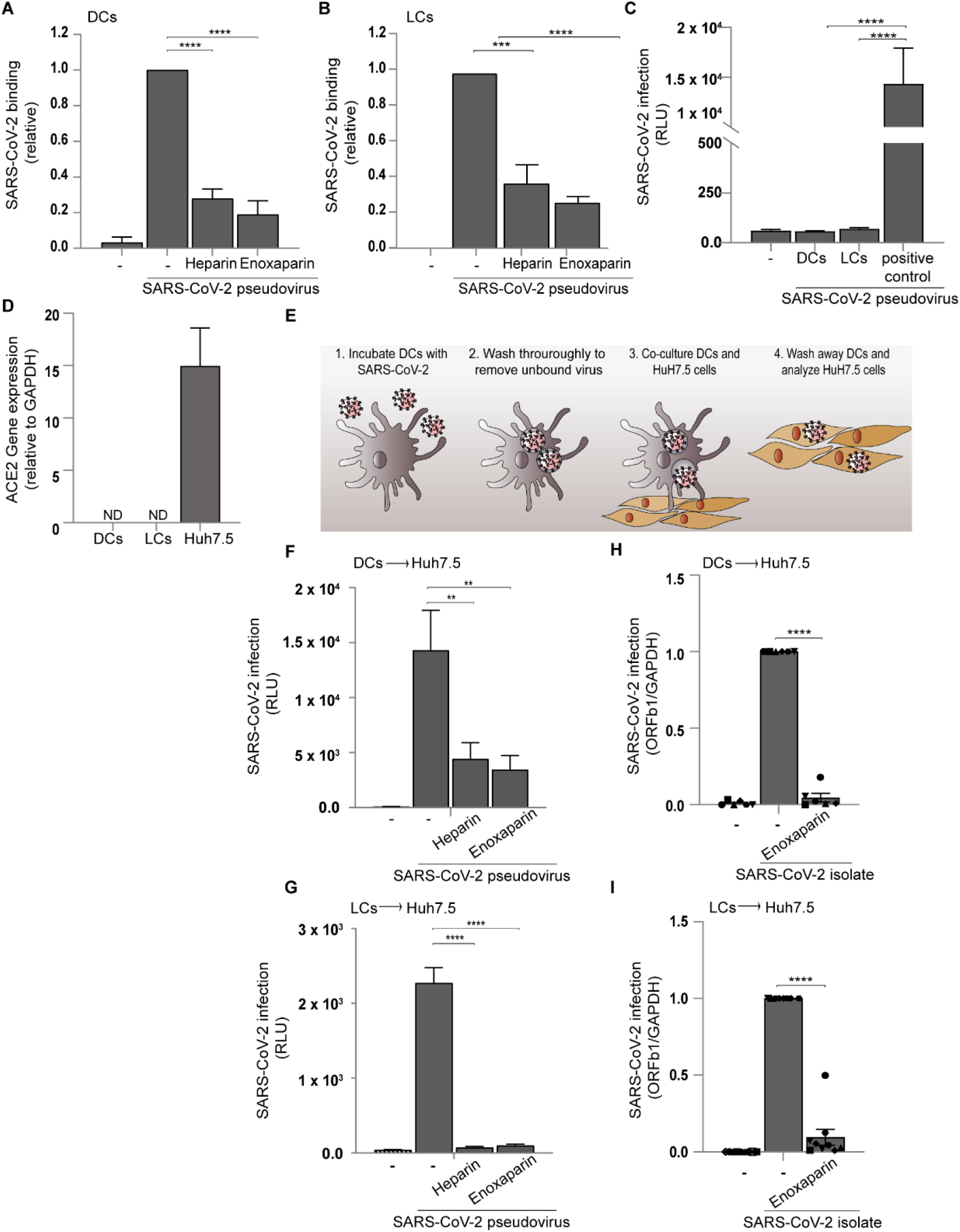
SARS-CoV-2 targets dendritic cells for dissemination. (A-B) SARS-CoV-2 binding to monocyte-derived DCs (A) and primary LCs (B) in presence of UF heparin (250IU) or enoxaparin (250IU). (C) DCs and LCs were infected with SARS-CoV-2 pseudovirus and infection was measured after 5 days by luciferase assay. As positive controls Huh 7.5 cells were infected.(D) ACE2 cell surface expression on DCs, LCs and Huh 7.5. Representative data for an experiment repeated more than three times with similar results. (E) Graphical overview of the cell-to-cell viral transmission. (F-G) DCs (F) and LCs (G) were preincubated with SARS-CoV-2 pseudovirus for 4 h in presence or absence of UF heparin (250 IU) or enoxaparin (250 IU), extensively washed and co-cultured with Huh 7.5 cells. Transmission by DCs or LCs to Huh 7.5 cells was determined by luciferase assay. (H-I) DCs (H) and LCs (I) were infected with SARS-CoV-2 isolate (Italy TCID50 104) for 24h in presence or absence of UF heparin (250IU) or enoxaparin (250IU), washed thoroughly and co-cultured with Huh7.5 cells. Quantification of viral RNA was measured by quantitative real-time PCR. Data show the mean values and error bars are the SEM. (A-B) ordinary one-way ANOVA with Tukey’s multiple-comparison test. (A)****p< 0.0001, (B) ***p= 0.0002, ****p< 0.0001 (n = 4), (C) ordinary one-way ANOVA with Tukey’s multiple-comparison test. ****p< 0.0001 (n=3 measured in triplicate), (F) ordinary one-way ANOVA with Dunnett’s multiple-comparison test. **p= 0.0042, **p= 0.0016 (n = 4 measured in triplicate), (G) ordinary one-way ANOVA with Tukey’s multiple-comparison test. ****p< 0.0001 (n = 3 measured in triplicate). (H-I) ordinary one-way ANOVA with Tukey’s multiple-comparison test. (H) ****p < 0.0001 (n=3 in duplicates), (I) ****p < 0.0001 (n=4 in triplicates). DCs: Dendritic cells, LCs: Langerhans cells, RLU: relative light units, ND: Not determined.

Different DC subsets transmit HIV-1 to target cells independent of productive infection (Geijtenbeek *et al*, 2000; Gurney *et al*, 2005; Sarrami-Forooshani *et al*., 2014). We therefore incubated DCs and LCs with SARS-CoV-2 pseudovirus and after washing co-cultured the DC subsets with susceptible Huh 7.5 cells (Fig 6E). Notably, co-culture of both virus-exposed DC subsets with Huh 7.5 cells led to infection, as determined by luciferase activity and infection was blocked by pre-treatment of pseudovirus with UF heparin and LMWH enoxaparin (Fig 6F and G). Next, DCs and LCs were incubated with the primary SARS-CoV-2 isolate and after extensive washing added to ACE-2 positive Huh 7.5 cells, and infection of Huh 7.5 cells was determined after removal of the DCs by quantitative qPCR. Co-culture of virus-exposed DCs and LCs with Huh 7.5 cells strongly increased infection, and transmission was inhibited by LMWH enoxaparin (Fig 6H and I). These data suggest that both DCs and LCs capture SARS-CoV-2 pseudovirus as well as the primary SARS-CoV-2 isolate via heparan sulfate proteoglycans and transmit the virus to ACE-2 expressing target cells.

### SARS-CoV-2 attaches to and infects primary nasal cells via heparan sulfate proteoglycans

Nasal epithelium is an important target for SARS-CoV-2 infection. Higher viral loads are detected in nasal and nasopharyngeal swabs compared to throat swabs (Tsang *et al*, 2021; Wang *et al*, 2020). Here we isolated nasal cells from healthy volunteers by brushing the inside of the nasal cavity. Epithelial cells are a major component of the isolated cells as shown by high percentage of cells positive for the epithelial cell marker EpCAM (Fig 7A). Also hematopoietic cells were present in the nasal cell fraction as shown by the expression of the hematopoietic cell marker CD45 (Fig 7A). Next we analyzed Syndecan 1 and 4 transcripts in the nasal fraction as well as expression of ACE-2. Especially high levels of Syndecan 1 transcripts were identified in the nasal fraction compared to those observed for polarized Calu-3 cells (Fig 7B). ACE2 transcript were also detected, suggesting that SARS-CoV-2 could directly infect nasal cells (Fig 7B) (Sungnak *et al*, 2020). We first investigated binding of primary SARS-CoV-2 isolate to the nasal cells. SARS-CoV-2 attached to the nasal cells from different donors and binding was blocked by LMWH enoxaparin (Figure 7C). Primary nasal cells were exposed to SARS-CoV-2 and cultured for 24 hours. Viability was not affected as measured by GAPDH expression. Notably, we observed high levels of SARS-CoV-2 ORF1b in cell-lysates (Figure 7D). Infection was inhibited by ACE2 block and, importantly, LMWH treatment blocked infection of SARS-CoV-2 as shown by decreased ORF1b in cell-lysate (Fig 7D). These data suggest that heparan sulfate proteoglycans expressed by the nasal epithelium are involved in SARS-CoV-2 binding and infection.

**Fig 7.**
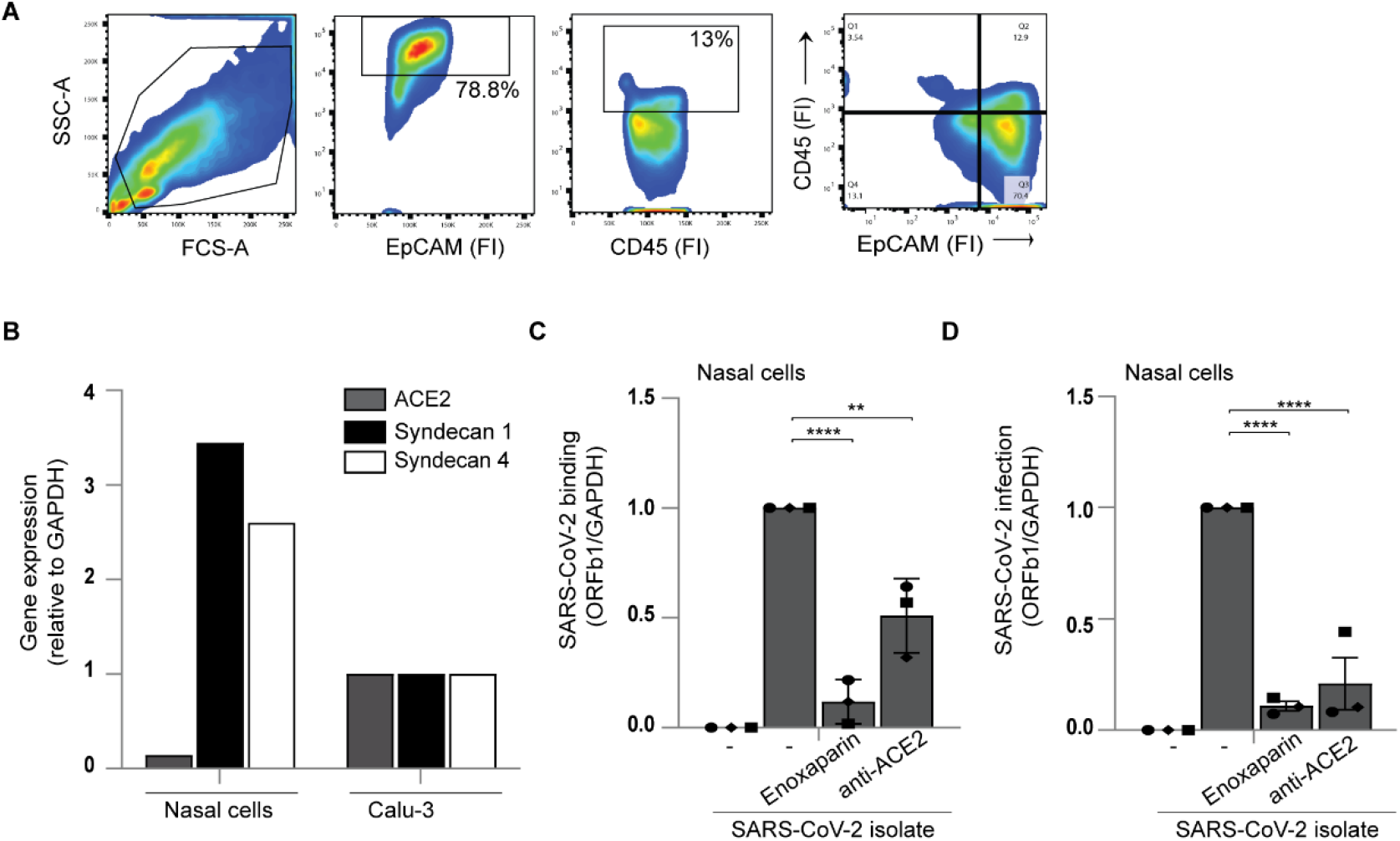
SARS-CoV-2 attaches to and infects primary nasal cells via heparan sulfate proteoglycans. (A) Flow cytometry analysis of single cell suspensions from the nasal epithelium. Cells were labelled with EpCAM and CD45 antibodies and gated accordingly. (B) ACE2, Syndecan 1 and Syndecan 4 cell surface expression on nasal cells, compared to polarized epithelial Calu-3 was confirmed by quantitative real-time PCR. (C-D) Nasal epithelial cells were exposed to a SARS-CoV-2 isolate (Italy TCID50 104) in presence of antibodies against ACE2 and enoxaparin (250IU). Detection of viral binding (C) and persistently-infected cells (D) was determined by quantitative real-time PCR. Data show the mean values and error bars are the SEM. Statistical analysis was performed using (C-D) ordinary on-way ANOVA with Tukey’s multiple-comparison test. (C) ****p < 0.0001, **p= 0.0013 (n=3), (D) ****p < 0.0001 (n=3 in duplicates),

## Discussion

SARS-CoV-2 interacts with ACE2 to infect cells. Recent studies suggest that heparan sulfates might interact with S protein to enhance viral attachment (Clausen *et al*., 2020; Zhang *et al*., 2020). Moreover, Clausen *et al*. show that heparan sulfate binding to SARS-CoV-2 facilitates ACE2 interactions (Clausen *et al*., 2020). Here we show that heparan sulfate proteoglycans on primary epithelial cells and primary dendritic cell subsets interact with both pseudotyped and primary SARS-CoV-2. We have identified Syndecan 1 and 4 as important attachment receptors for SARS-CoV-2. Interestingly, neutralizing antibodies against SARS-CoV-2 prevented the interaction of SARS-CoV-2 with Syndecan 1, suggesting that antibodies targeting the interaction of SARS-CoV-2 with heparan sulfates might also neutralize infection as was shown for antibodies against ACE2. Moreover, we identified heparan sulfate proteoglycans as important attachment receptors facilitating transmission by primary mucosal DC subsets independent of infection. Both UF heparin and LWMH efficiently blocked infection and transmission of SARS-CoV-2. Moreover, we show that LMWH efficiently block infection of primary nasal epithelial cells. Thus, heparan sulfate proteoglycans function as attachment receptors for SARS-CoV-2 on primary epithelial and dendritic cells, and targeting these receptors might prevent infection.

Our data indicate that SARS-CoV-2 binding to polarized colorectal and respiratory epithelial cells is dependent on heparan sulfates, supporting a role for heparan sulfates as attachment receptors. Moreover, infection of polarized respiratory epithelial cells by SARS CoV-2 hCoV-19/Italy strain as well as pseudovirus was inhibited by LMWH to a similar level as anti-ACE2 antibodies. Combinations of LMWH with antibodies did not further decrease infection. These data suggest that SARS-CoV-2 attaches to cells via heparan sulfate proteoglycan, which facilitates interaction with ACE2 and subsequent infection.

Neutralizing antibodies against SARS-CoV-2 are a potential therapy for COVID-19 patients and most potent monoclonal neutralizing antibodies target the RBD site of the S protein thereby preventing interaction of S protein with ACE2 (Brouwer *et al*., 2020). However, neutralization antibodies have also been isolated that target non-RBD sites of the S protein. Indeed, COVA1-21 targets a non-RBD site as it does not seem to interfere with ACE2 (Brouwer *et al*., 2020), suggesting that it neutralizes either via another receptor or another mechanism. We screened different antibodies isolated from COVID-19 patients (Brouwer *et al*., 2020) for blocking SARS-CoV-2 binding to Syndecan 1. Two RBD antibodies COVA1-18 and COVA2-15 and one non-RBD antibody COVA1-21 were identified that blocked interaction of Syndecan 1 with SARS-CoV-2 pseudovirus as well as the SARS-CoV-2 hCoV-19/Italy strain. Notably, these three antibodies are potent neutralizing antibodies against SARS-CoV-2 (Brouwer *et al*., 2020). Most non-neutralizing antibodies did not interfere with SARS-CoV-2 binding to Syndecan 1. These data suggest that blocking the interaction of SARS-CoV-2 with heparan sulfate proteoglycans might be a new mechanism of neutralization.

Different DC subsets are present in mucosal tissues to capture pathogens for antigen presentation. After pathogen interactions, DCs migrate into lymphoid tissues (Randolph *et al*, 2005). Several viruses such as HIV-1 and Dengue virus, hijack DC functions for dissemination (Geijtenbeek *et al*., 2000; Pham *et al*, 2012). Primary LCs and DCs efficiently captured SARS-CoV-2 via heparan sulfate proteoglycans. Previously we have shown that LCs express Syndecan 4 (Nijmeijer *et al*., 2020) and DCs express Syndecan 3 and Syndecan 4 (de Witte *et al*, 2007a). Thus, our data suggest that both Syndecan 3 and 4 might be involved in SARS-CoV-2 capture. Both LCs and DCs did not express ACE2 and were not infected by SARS-CoV-2 pseudovirus. However, co-culture of SARS-CoV-2-exposed LCs and DCs with ACE2 cells led to productive infection with pseudotyped as well as a primary SARS-CoV-2 strain. Transmission was blocked by LMWH suggesting that capture by heparan sulfate proteoglycans is important for transmission. Moreover, our data indicate that SARS-CoV-2 transmission is independent of direct infection of DC subsets, suggesting that this might allow the virus from escaping neutralization by antibodies or antiviral drugs.

The upper airways and nasal epithelium might be the primary route of infection as higher viral load have been found in nasal swabs when compared to throat swabs (Zou *et al*, 2020a) Moreover, nasal epithelial cells express angiotensin-converting enzyme 2 (ACE2) and the cellular serine protease TMPRSS2 (Sungnak *et al*., 2020). We have isolated nasal cells from healthy volunteers using a nasal brush and the majority of cells were EpCAM-positive epithelial cells and some hematopoietic cells, most likely lymphocytes and myeloid cells. Syndecan 1 and 4 transcripts were detected at high levels in nasal cell fraction, suggesting that Syndecans might be involved in virus interactions. Our data support an important role for nasal cells as the first target for SARS-CoV-2 as nasal cells efficiently captured primary SARS-CoV-2 and were also infected by SARS-CoV-2. LMWH blocked capture and infection of the nasal cells. Interestingly, the nasal cells were not cultured as done in previous studies (Müller *et al*, 2013; Vanders *et al*, 2019) suggesting that the nasal epithelial cells are a direct target for SARS-CoV-2 and that heparan sulfate proteoglycans are involved in the infection.

LMWHs are already used as subcutaneous treatment of COVID-19 patients to prevent systemic clotting (World Health Organization, 2020a; Zhai *et al*, 2020). Interestingly, here we have identified an important ability of LMWH to directly block SARS-CoV-2 binding and infection of epithelial cells as well as preventing virus transmission. Our data support the use of LMWH as prophylactic treatment for SARS-CoV-2 as well as a treatment option early in infection to block further infection and dissemination. Vaccination programs are currently running worldwide but it remains unclear whether this is sufficient for specific patients who are immunocompromised or suffer from other diseases that prevent an efficient immune response upon vaccination. LMWH prophylaxis might also be used when new SARS-CoV-2 variants arise that are not efficiently counteracted by the current vaccines.

## Materials and Methods

### Reagents and antibodies

The following antibodies were used (all anti-human): ACE-2 (R&D), (Heparan Sulfate (clone F58-10E4) (Amsbio), digested Heparan (clone F69-3G10) (Amsbio), CD1a-APC mouse IgG1 (BD Biosciences, San Jose, CA, USA), CD207-PE (langerin) mouse IgG1 (#IM3577), PerCP-Cy™ 5.5-conjugated mouse IgG1 EPCAM 347199) (BD Bioscience), PE-conjugated mouse IgG1 E-Cadherin (FAB18381P) (R&D Systems), APCcy-conjugated mouse IgG1 CD45 (557833) (BD Bioscience), APC-conjugated CD14 (21620146sp) (Immunotools), PE-conjugated mouse IgG1 CD11b (101208) (Biolegend). FITC-conjugated goat-anti-mouse IgM (#31992) (Invitrogen), AF488-conjugated donkey-anti-mouse IgG2b (Invitrogen). Flow cytometric analyses were performed on a BD FACS Canto II (BD Biosciences). Data was analyzed using FlowJo vX.0.7 software (TreeStar).

The following reagents were used: Unfractionated (UF) heparin, 5.000 I.E./ml (LEO). Low Molecular Weight heparins (LMWH): dalteparin, 10.000 IE anti-Xa/ml (Pfizer), tinzaparin, 10.000 IE anti-X1/0.5ml (LEO), enoxaparin, 6000 IE (60mg)/0.6 ml (Sanofi), nadroparin, 9.500 IE anti-XA/ml (Aspen). Heparinase III from *Flavobacterium heparium*, EC 4.2.2.8, Batch 010, (Amsbio). Biotinylated SARS-CoV-2 S protein as well as neutralizing and non-neutralizing COVA antibodies were generated as described previously (Brouwer *et al*., 2020).

### Cell lines

The Simian kidney cell line VeroE6 (ATCC® CRL-1586™) was maintained in CO2 independent medium (Gibco Life Technologies, Gaithersburg, Md.) supplemented with 10% fetal calf serum (FCS), L-glutamine and penicillin/streptomycin (10 μg/ml). Culture was maintained at 37C without CO2. Huh7.5 (human hepatocellular carcinoma) cells received from dr. Charles M. Rice (Lindenbach *et al*, 2005) were maintained in Dulbecco modified Eagle medium (Gibco Life Technologies, Gaithersburg, Md.) containing 10% fetal calf serum (FCS), L-glutamine and penicillin/streptomycin (10 μg/ml). Medium was supplemented with 1mM Hepes buffer (Gibco Life Technologies, Gaithersburg, Md.). The human B cell line Namalwa (ATCC, CRL-1432) and Namalwa cells stably expressing human Syndecan 1 and Syndecan 4 (Zhang *et al*, 2001) were a gift from Dr. Guido David and Dr. Philippe A Gallay. The cells were maintained in RPMI 1640 medium (Gibco Life Technologies, Gaithersburg, Md.) containing 10% fetal calf serum (FCS), penicillin/streptomycin (10 μg/ml) and 1 mM sodium pyruvate (Thermo Fisher). The expression of the different Syndecans was validated by PCR analysis using specific primers aimed against Syndecans. The human multiple myeloma cell line XG-1 was cultured in Iscove modified Dulbecco medium (Invitrogen Life Technologies) containing 10% fetal bovine serum, 100 U/mL of penicillin and 100 µg/mL of streptomycin. The medium was further was supplemented with 500 pg/mL of interleukin-6 (Prospec). The CRISPR-Cas9 knockout for *Ext1* has been described previously (Ren *et al*., 2018). The human embryonic kidney 293T/17 cells (ATCC, CRL-11268) were maintained in maintained in Dulbecco modified Eagle medium (Gibco Life Technologies, Gaithersburg, Md.) containing 10% fetal calf serum (FCS), L-glutamine and penicillin/streptomycin (10 μg/ml). The human epithelial Caco-2 cells (ATCC, HTB-37™) as well as the human lung epithelial Calu-3 cells (ATCC® HTB-55™) were maintained in Dulbecco modified Eagle medium (Gibco Life Technologies, Gaithersburg, Md.) containing 10% fetal calf serum (FCS), L-glutamine and penicillin/streptomycin (10 μg/ml) and supplemented with MEM Non-Essential Amino Acids Solution (NEAA) (Gibco Life Technologies, Gaithersburg, Md.). To create a monolayer of polarized cells, Caco-2 and Calu-3 cells were maintained in 6.5 mm Transwell® with 5.0 µm Pore Polycarbonate Membrane Insert (Corning). The cells were initially seeded with a density of 25.000 cells per 6.5 mm filter insert and full polarization was reached after 2 weeks in culture. Polarization was monitored by measuring transepithelial electrical resistance (TEER).

### Primary human cells

This study has been conducted in accordance with the ethical principles set out in the declaration of Helsinki and was approved by the institutional review board of the Academic Medical Center (AMC, Amsterdam, Netherlands) and the Ethics Advisory Body of the Sanquin Blood Supply Foundation (Amsterdam, Netherlands).

CD14+ monocytes were isolated from the blood of healthy volunteer donors (Sanquin blood bank) and subsequently differentiated into monocyte-derived DCs as described previously (Mesman *et al*, 2014) (Fig EV3b). LCs were isolated from human epidermal sheets obtained from healthy donors after plastic surgery. Epidermal sheets were prepared as described previously (de Witte *et al*., 2007b; Sarrami-Forooshani *et al*., 2014). Briefly, skin-grafts were obtained using a dermatome (Zimmer Biomet, Indiana USA). After incubation with Dispase II (1 U/ml, Roche Diagnostics), epidermal sheets were separated from dermis, washed and cultured in IMDM (Thermo Fischer Scientific, USA) supplemented with 10% FCS, gentamycine (20 μg/ml, Centrafarm, Netherlands), pencilline/streptomycin (10 U/ml and 10 μg/ml, respectively; Invitrogen) for 3 days after which LCs were harvested. Purity of LCs was routinely verified by flow cytometry using antibodies directed against CD207 (langerin) and CD1a (Fig EV3A).

Primary nasal epithelial cells were obtained from healthy volunteers. Cells were isolated from the lower nasal cavity with a brush after which they were transferred into CO2 independent medium (Gibco Life Technologies, Gaithersburg, Md.) supplemented with 10% fetal calf serum (FCS), L-glutamine and penicillin/streptomycin (10 μg/ml). Cell surface receptor expression was determined by flow cytometer.

### SARS-Cov-2 pseudovirus production

For production of single-round infection viruses, human embryonic kidney 293T/17 cells (ATCC, CRL-11268) were co-transfected with an adjusted HIV backbone plasmid (pNL4-3.Luc.R-S-) containing previously described stabilizing mutations in the capsid protein (PMID: 12547912) and firefly luciferase in the *nef* open reading frame (1.35ug) and pSARS-CoV-2 expressing SARS-CoV-2 S protein (0.6ug) (GenBank; MN908947.3) (Brouwer *et al*., 2020).Transfection was performed in 293T/17 cells using genejuice (Novagen, USA) transfection kit according to manufacturer’s protocol. At day 3 or day 4, pseudotyped SARS-CoV-2 virus particles were harvested and filtered over a 0.45 µm nitrocellulose membrane (SartoriusStedim, Gottingen, Germany). SARS-CoV-2 pseudovirus productions were quantified by p24 ELISA (Perkin Elmer Life Sciences).

### SARS-CoV-2 WT virus production

The following reagent was obtained from Dr. Maria R. Capobianchi through BEI Resources, NIAID, NIH: SARS-Related Coronavirus 2, Isolate Italy-INMI1, NR-52284, originally isolated January 2020 in Rome, Italy. VeroE6 cells (ATCC® CRL-1586™) were inoculated with the SARS-CoV-2 isolate and used for reproduction of virus stocks. CPE formation was closely monitored and virus supernatant was harvested after 48 hours. Tissue culture infectious dose (TCID50) was determined on VeroE6 cells by MTT assay 48 hours after infection. Loss of MTT staining as determined by spectrometer is indicative of cell death.

### Pseudovirus infection assays

HuH7.5 and 293T(+hACE2) cells were exposed to 95 ng of pseudotyped SARS-CoV-

2. Primary dendritic cell subsets were exposed to 190 ng and polarized Caco2 and Calu3 cells to 477.62 ng of pseudotyped SARS-CoV-2. Virus was pre-incubated with 250U LMWH or UF heparin prior to addition of cells. Infection was measured after 5 days at 37°C by the Luciferase assay system (Promega, USA) according to manufacturer’s instructions.

### SARS-CoV-2 WT infection assays

VeroE6 cells were exposed to the SARS-CoV-2 isolate (hCoV-19/Italy) at different TCIDs for 48 hours. Virus was pre-incubated with 250IU of LMWH prior to addition to the cells. Infection was measured after 48 hours at 37°C by MTT and determined by cell viability.

### Tetrazolium dye (MTT) colorimetric cell viability assay

MTT solution was added to VeroE6 cells and incubated for 2 hours at 37°C. After removing the MTT solution, MTT solvent containing 4 mM HCL and 1% Nonidet P-40 (NP40) in isopropanol was added to the cells. Homogenous solution was measured at optical density between 580 nm and 655 nm.

### 293T Transfection with ACE2

To generate cells expressing human ACE2, human embryonic kidney 293T/17 cells were transfected with pcDNA3.1(-)hACE2 (Addgene plasmid #1786). Transfection was performed in 293T/17 cells using the genejuice (Novagen, USA) transfection kit according to manufacturer’s protocol. At 24h post-transfection, cells were washed with phosphate-buffered saline (PBS) and cultured for recovering at 37C for 24h in Dulbecco’s MEM supplemented with 10% heat-inactivated fetal calf serum (FCS), L-glutamine and penicillin/streptomycin (10 U/ml) After 24h of recovery, cells were cultured in media supplemented with G418 (5mg/mL) (Thermo Fisher) and passage for 3 weeks at 37C. Surviving clones were analyzed for ACE2 expression via flow cytometry and PCR.

### Virus binding and sensitive p24 ELISA

In order to determine SARS-CoV-2 binding, target cells were exposed to 95 ng of pseudotyped SARS-CoV-2 virus for 4 hours at 4°C. Cells were washed to remove the unbound virus and lysed with lysis buffer. Binding and internalization were quantified by RETRO-TEK HIV-1 p24 ELISA according to manufacturer instructions (ZeptoMetrix Corporation).

### Transmission assays and co-culture

DCs or LCs were exposed to 191.05 ng of pseudotyped SARS-CoV-2 or pseudotyped SARS-CoV-2 pre-incubated with 250U UF heparin or LMWH for 4 hours, harvested, extensively washed to remove unbound virus and co-cultured with Huh7.5 for 5 days at 37°C after which they were analyzed for with the Luciferase assay system (Promega, USA) according to manufacturer’s instructions. DCs or LCs were further exposed to 10 TCID or SARS-CoV-2 isolate or SARS-Cov-2 isolated pre-incubated with 250 IU LMWH for 24 hours, extensively washed to remove unbound virus and co-cultured with HuH7.5 for 24 hours at 37°C. After 24 hours, HuH7.5 were again washed extensively to remove DCs or LCs and lysed for isolation of viral RNA. DC markers (dectin-1, langerin) were analyzed to determine the efficiency of the washing steps and >90% DCs were removed after washing.

### RNA isolation and quantitative real time PCR

RNA of cells exposed to SARS-CoV-2 was isolated with the QIAamp Viral RNA Mini Kit (Qiagen) according to the manufacturers protocol. cDNA was synthesized with the M-MLV reverse-transcriptase kit (Promega) and diluted 1 in 5 before further application. Cellular mRNA of cells not exposed to virus was isolated with an mRNA Capture kit (Roche) and cDNA was synthesized with a reverse-transcriptase kit (Promega). PCR amplification was performed in the presence of SYBR green in a 7500 Fast Realtime PCR System (ABI). Specific primers were designed with Primer Express 2.0 (Applied Biosystems). Primer sequences used for mRNA expression were for gene product: GAPDH, forward primer (CCATGTTCGTCATGGGTGTG), reverse primer (GGTGCTAA GCAGTTGGTGGTG). For gene product SARS-CoV-2 ORf1b, forward primer (TGGGGTTTTACAGGTAACCT), reverse primer (AACACGCTTAAC AAAGCACTC) as described previously (Chu *et al*, 2020). For gene product: ACE2, forward primer (GGACCCAGGAAATGTTCAGA), reverse primer (GGCTGCAGAAA GTGACATGA). For gene product: Syndecan 1, forward primer (ATCACCTTGTCACAGCAGACCC) reverse primer (CTCCACTTCTGG CAGGACTACA). Syndecan 4, forward primer (AGGTGTCAATGTCCAGCACTGTG) reverse primer (AGCAGTAGGATCAGGAAGACGGC). The normalized amount of target mRNA was calculated from the Ct values obtained for both target and household mRNA with the equation Nt = 2Ct (GAPDH) − Ct(target). For relative mRNA expression, control siRNA sample was set at 1 for each donor.

### Biosynthesis inhibition and enzymatic treatment

HuH7.5 cells were treated in D-PBS/0.25% BSA with 46 miliunits heparinase III (Amsbio) for 1 hour at 37°C, washed and used in subsequent experiments. Enzymatic digestion was verified by flow cytometry using antibodies directed against heparan sulfates and digested heparan sulfates.

### Statistics

All results are presented as mean ± SEM and were analyzed by GraphPad Prism 8 software (GraphPad Software Inc.). A two-tailed, parametric Student’s *t*-test for paired observations (differences within the same donor) or unpaired observation, Mann-Whitney tests (differences between different donors, that were not normally distributed) was performed. For unpaired, non-parametric observations a one-way ANOVA or two-way ANOVA test with post hoc analysis (Tukey’s or Dunnet’s) were performed. Statistical significance was set at *P< 0.05, **P<0.01***P<0.001****P<0.0001.

## Acknowledgements

We thank Jonne Snitselaar and Yoann Aldon for help with production of antibodies and Hildo Lantermans for help with the XG1/EXT1 KO cell lines.

## Author Contributions

M.B-J and J.E conceived and designed experiments; M.B-J, J.E, T.M.K, L.C.H, J.L.v.H performed the experiments and contributed to scientific discussion; P.J.M.B., K.E.V, A.P.J. V, F.E.H.P.v.B, M.S, G.J.d.B, R.W.S., M.J.v.G, B.M.N and N.A.K contributed essential research materials and scientific input. M.B-J, J.E, T.M.K and T.B.H.G analyzed and interpreted data; J.E, M.B-J and T.B.H.G. wrote the manuscript with input from all listed authors. T.B.H.G. was involved in all aspects of the study.

## Supplementary material

**Fig EV1.**
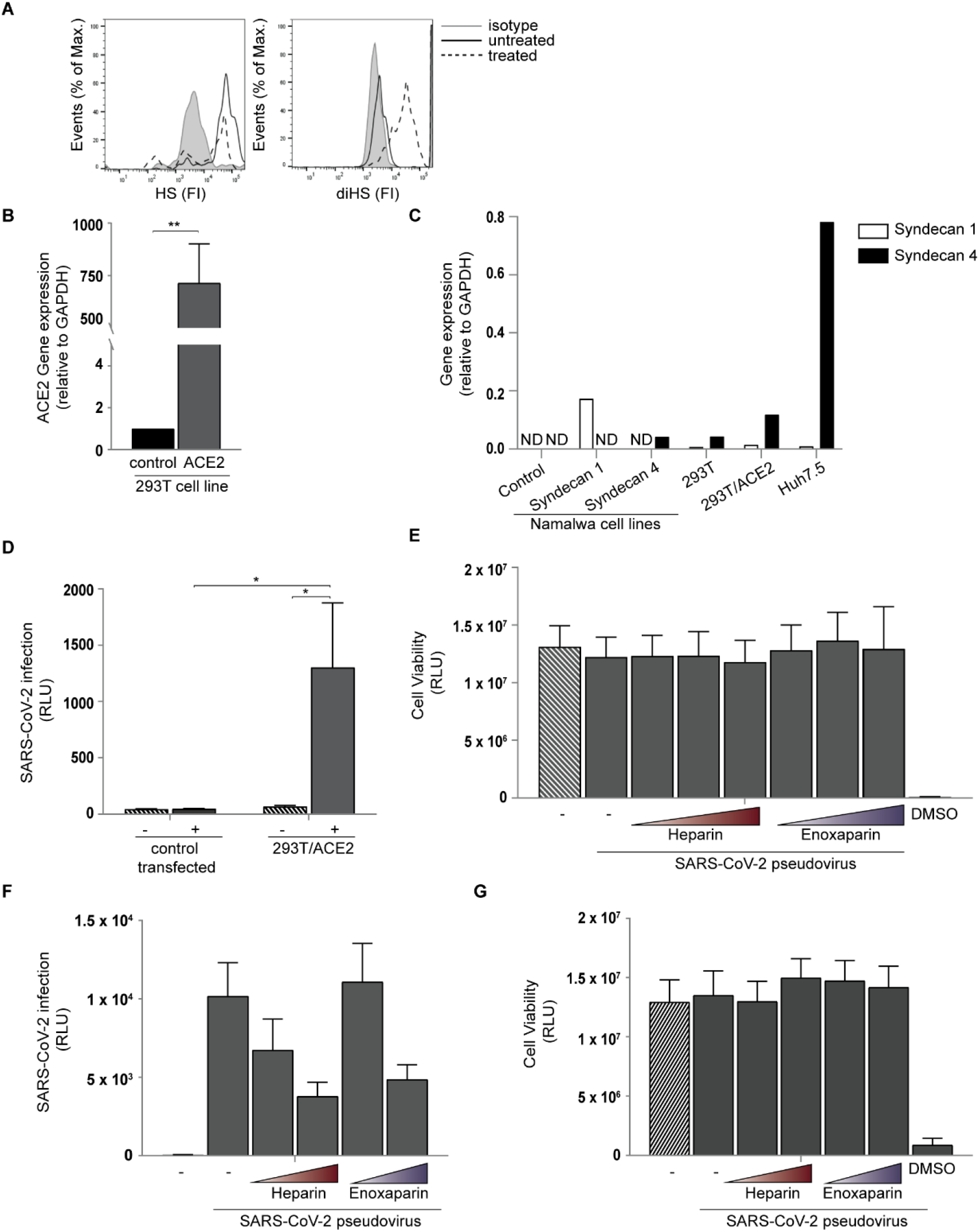
(A) Huh7.5 were left untreated or treated with heparinase for 1 h and heparan sulfate or digested heparan sulfate expression was determined by flow cytometry. One representative donor out of 3 is depicted. (B) Cell surface expression of ACE2 on 293T (control and ACE2 transfected) was determined by quantitative real-time PCR. (C) Syndecan 1 and Syndecan 4 cell surface expression on Namalwa cell lines, 293T cell lines and Huh7.5 was confirmed by quantitative real-time PCR. Representative data for an experiment repeated more than three times with similar results. (D) SARS-CoV-2 pseudovirus infection on 293T (control vs ACE2-transfected cells) was measured by luciferase assay. (E) Cell Viability of infected Huh7.5 with SARS-CoV-2 pseudovirus in presence of different concentration of UF heparin and LMWH enoxaparin. (F-G) VeroE6 cells were infected with a heparin and enoxaparin pretreated SARS-CoV-2 pseudovirus. Infection (F) and cell viability (G) was determined by luciferase assay. Data show the mean values and error bars are the SEM. Statistical analysis was performed using (C) 2way-ANOVA with Dunnett’s multiple-comparison test. *p=0.0198, *p=0.0206 (n=3 in triplicates), (F) ordinary one-way ANOVA with Tukey’s multiple-comparison test. RLU: relative light units, ND: Not determined.

**Fig EV2.**
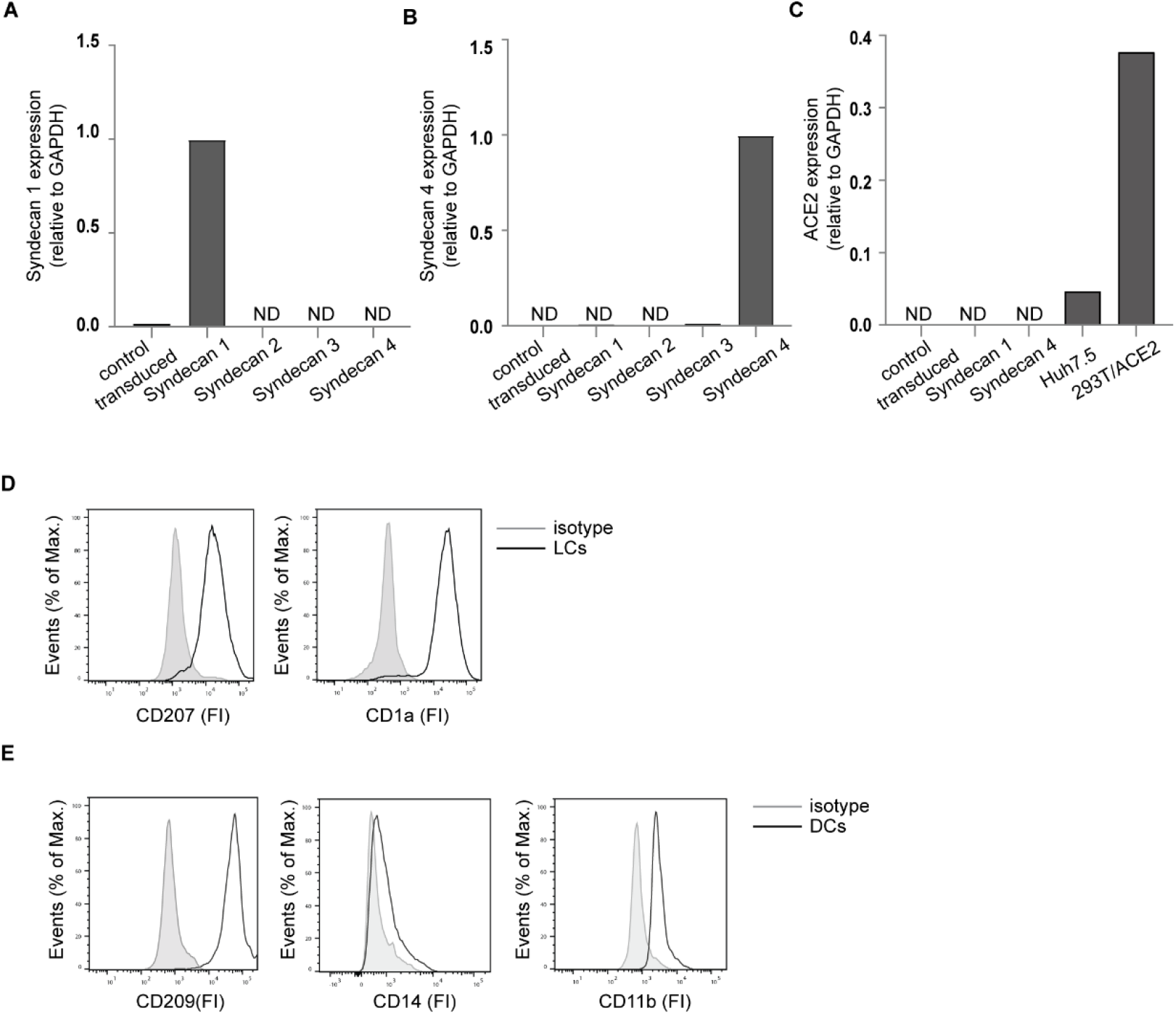
(A-B) Different Namalwa Syndecan cell lines express Syndecan 1 (A) – 4 (B), on the cell surface determined by quantitative real-time PCR. (C) ACE2 expression was confirmed for different Namalwa Syndecan cell lines in comparison to Huh7.5 cell line and ACE2-transfected 293T. (D) LCs were stained with antibodies against CD207 and CD1a and analysed by flow cytometry. The histogram shows the cell surface expression of the receptor. (E) DCs were stained with antibodies against the surface markers CD209, CD14 and CD11b and analysed by flow cytometry. DCs: Dendritic cells, LCs: Langerhans cells, RLU: relative light units, ND: Not determined.

## References

Agha M, Blake M, Chilleo C, Wells A, Haidar G (2021) Suboptimal response to COVID- 19 mRNA vaccines in hematologic malignancies patients. medRxiv

Artursson P, Palm K, Luthman K (2001) Caco-2 monolayers in experimental and theoretical predictions of drug transport. Adv Drug Deliv Rev 46: 27–43

Bacsa S, Karasneh G, Dosa S, Liu J, Valyi-Nagy T, Shukla D (2011) Syndecan-1 and syndecan-2 play key roles in herpes simplex virus type-1 infection. J Gen Virol 92: 733–743

Bosch BJ, van der Zee R, de Haan CA, Rottier PJ (2003) The coronavirus spike protein is a class I virus fusion protein: structural and functional characterization of the fusion core complex. J Virol 77: 8801–8811

Boyarsky BJ, Ruddy JA, Connolly CM, Ou MT, Werbel WA, Garonzik-Wang JM, Segev DL, Paik JJ (2021) Antibody response to a single dose of SARS-CoV-2 mRNA vaccine in patients with rheumatic and musculoskeletal diseases. Ann Rheum Dis

Brooks SK, Webster RK, Smith LE, Woodland L, Wessely S, Greenberg N, Rubin GJ (2020) The psychological impact of quarantine and how to reduce it: rapid review of the evidence. Lancet 395: 912–920

Brouwer PJM, Caniels TG, van der Straten K, Snitselaar JL, Aldon Y, Bangaru S, Torres JL, Okba NMA, Claireaux M, Kerster G et al (2020) Potent neutralizing antibodies from COVID-19 patients define multiple targets of vulnerability. Science 369: 643–650

Burkard C, Verheije MH, Wicht O, van Kasteren SI, van Kuppeveld FJ, Haagmans BL, Pelkmans L, Rottier PJ, Bosch BJ, de Haan CA (2014) Coronavirus cell entry occurs through the endo-/lysosomal pathway in a proteolysis-dependent manner. PLoS Pathog 10: e1004502

Byrnes AP, Griffin DE (1998) Binding of Sindbis virus to cell surface heparan sulfate. J Virol 72: 7349–7356

Chu DKW, Pan Y, Cheng SMS, Hui KPY, Krishnan P, Liu Y, Ng DYM, Wan CKC, Yang P, Wang Q et al (2020) Molecular Diagnosis of a Novel Coronavirus (2019-nCoV) Causing an Outbreak of Pneumonia. Clin Chem 66: 549–555

Clausen TM, Sandoval DR, Spliid CB, Pihl J, Perrett HR, Painter CD, Narayanan A, Majowicz SA, Kwong EM, McVicar RN et al (2020) SARS-CoV-2 Infection Depends on Cellular Heparan Sulfate and ACE2. Cell 183: 1043–1057.e1015

Collier DA, De Marco A, Ferreira I, Meng B, Datir RP, Walls AC, Kemp SA, Bassi J, Pinto D, Silacci-Fregni C et al (2021) Sensitivity of SARS-CoV-2 B.1.1.7 to mRNA vaccine-elicited antibodies. Nature 593: 136–141

de Witte L, Bobardt M, Chatterji U, Degeest G, David G, Geijtenbeek TB, Gallay P (2007a) Syndecan-3 is a dendritic cell-specific attachment receptor for HIV-1. Proc Natl Acad Sci U S A 104: 19464–19469

de Witte L, Nabatov A, Pion M, Fluitsma D, de Jong MA, de Gruijl T, Piguet V, van Kooyk Y, Geijtenbeek TB (2007b) Langerin is a natural barrier to HIV-1 transmission by Langerhans cells. Nat Med 13: 367–371

Ferioli M, Cisternino C, Leo V, Pisani L, Palange P, Nava S (2020) Protecting healthcare workers from SARS-CoV-2 infection: practical indications. Eur Respir Rev 29

Geijtenbeek TB, Kwon DS, Torensma R, van Vliet SJ, van Duijnhoven GC, Middel J, Cornelissen IL, Nottet HS, KewalRamani VN, Littman DR et al (2000) DC-SIGN, a dendritic cell-specific HIV-1-binding protein that enhances trans-infection of T cells. Cell 100: 587–597

Gurney KB, Elliott J, Nassanian H, Song C, Soilleux E, McGowan I, Anton PA, Lee B (2005) Binding and transfer of human immunodeficiency virus by DC-SIGN+ cells in human rectal mucosa. J Virol 79: 5762–5773

Hamming I, Timens W, Bulthuis ML, Lely AT, Navis G, van Goor H (2004) Tissue distribution of ACE2 protein, the functional receptor for SARS coronavirus. A first step in understanding SARS pathogenesis. J Pathol 203: 631–637

Harapan H, Itoh N, Yufika A, Winardi W, Keam S, Te H, Megawati D, Hayati Z, Wagner AL, Mudatsir M (2020) Coronavirus disease 2019 (COVID-19): A literature review. J Infect Public Health 13: 667–673

Harcourt JL, Caidi H, Anderson LJ, Haynes LM (2011) Evaluation of the Calu-3 cell line as a model of in vitro respiratory syncytial virus infection. J Virol Methods 174: 144–149

Hayashida K, Johnston DR, Goldberger O, Park PW (2006) Syndecan-1 expression in epithelial cells is induced by transforming growth factor beta through a PKA-dependent pathway. J Biol Chem 281: 24365–24374

Hoffmann M, Kleine-Weber H, Schroeder S, Krüger N, Herrler T, Erichsen S, Schiergens TS, Herrler G, Wu NH, Nitsche A et al (2020) SARS-CoV-2 Cell Entry Depends on ACE2 and TMPRSS2 and Is Blocked by a Clinically Proven Protease Inhibitor. Cell 181: 271–280.e278

Hui KPY, Cheung MC, Perera R, Ng KC, Bui CHT, Ho JCW, Ng MMT, Kuok DIT, Shih KC, Tsao SW et al (2020) Tropism, replication competence, and innate immune responses of the coronavirus SARS-CoV-2 in human respiratory tract and conjunctiva: an analysis in ex-vivo and in-vitro cultures. Lancet Respir Med

Hulswit RJ, de Haan CA, Bosch BJ (2016) Coronavirus Spike Protein and Tropism Changes. Adv Virus Res 96: 29–57

Jiang J, Cun W, Wu X, Shi Q, Tang H, Luo G (2012) Hepatitis C virus attachment mediated by apolipoprotein E binding to cell surface heparan sulfate. J Virol 86: 7256–7267

Jones KS, Petrow-Sadowski C, Huang YK, Bertolette DC, Ruscetti FW (2008) Cell-free HTLV-1 infects dendritic cells leading to transmission and transformation of CD4(+) T cells. Nat Med 14: 429–436

Kakkar AK (2004) Low- and ultra-low-molecular-weight heparins. Best Pract Res Clin Haematol 17: 77–87

Lamers MM, Beumer J, van der Vaart J, Knoops K, Puschhof J, Breugem TI, Ravelli RBG, Paul van Schayck J, Mykytyn AZ, Duimel HQ et al (2020) SARS-CoV-2 productively infects human gut enterocytes. Science

Letko M, Marzi A, Munster V (2020) Functional assessment of cell entry and receptor usage for SARS-CoV-2 and other lineage B betacoronaviruses. Nat Microbiol 5: 562–569

Li F, Li W, Farzan M, Harrison SC (2005) Structure of SARS coronavirus spike receptor-binding domain complexed with receptor. Science 309: 1864–1868

Lindenbach BD, Evans MJ, Syder AJ, Wölk B, Tellinghuisen TL, Liu CC, Maruyama T, Hynes RO, Burton DR, McKeating JA et al (2005) Complete replication of hepatitis C virus in cell culture. Science 309: 623–626

Marzi A, Gramberg T, Simmons G, Möller P, Rennekamp AJ, Krumbiegel M, Geier M, Eisemann J, Turza N, Saunier B et al (2004) DC-SIGN and DC-SIGNR interact with the glycoprotein of Marburg virus and the S protein of severe acute respiratory syndrome coronavirus. J Virol 78: 12090–12095

Mathieu E, Ritchie H, Ortiz-Ospina E, Roser M, Hasell J, Appel C, Giattino C, Rodés- Guirao L (2021) A global database of COVID-19 vaccinations. Nat Hum Behav

Merad M, Ginhoux F, Collin M (2008) Origin, homeostasis and function of Langerhans cells and other langerin-expressing dendritic cells. Nat Rev Immunol 8: 935–947

Mesman AW, Zijlstra-Willems EM, Kaptein TM, de Swart RL, Davis ME, Ludlow M, Duprex WP, Gack MU, Gringhuis SI, Geijtenbeek TB (2014) Measles virus suppresses RIG-I-like receptor activation in dendritic cells via DC-SIGN-mediated inhibition of PP1 phosphatases. Cell Host Microbe 16: 31–42

Milewska A, Zarebski M, Nowak P, Stozek K, Potempa J, Pyrc K (2014) Human coronavirus NL63 utilizes heparan sulfate proteoglycans for attachment to target cells. J Virol 88: 13221–13230

Müller L, Brighton LE, Carson JL, Fischer WA, 2nd, Jaspers I (2013) Culturing of human nasal epithelial cells at the air liquid interface. J Vis Exp

Nicola M, Alsafi Z, Sohrabi C, Kerwan A, Al-Jabir A, Iosifidis C, Agha M, Agha R (2020) The socio-economic implications of the coronavirus pandemic (COVID-19): A review. Int J Surg 78: 185–193

Nijmeijer BM, Eder J, Langedijk CJM, Kaptein TM, Meeussen S, Zimmermann P, Ribeiro CMS, Geijtenbeek TBH (2020) Syndecan 4 Upregulation on Activated Langerhans Cells Counteracts Langerin Restriction to Facilitate Hepatitis C Virus Transmission. Front Immunol 11: 503

Nijmeijer BM, Sarrami-Forooshani R, Steba GS, Schreurs RR, Koekkoek SM, Molenkamp R, Schinkel J, Reiss P, Siegenbeek van Heukelom ML, van der Valk M et al (2019) HIV-1 exposure and immune activation enhance sexual transmission of Hepatitis C virus by primary Langerhans cells. J Int AIDS Soc 22: e25268

Peiris JS, Yuen KY, Osterhaus AD, Stöhr K (2003) The severe acute respiratory syndrome. N Engl J Med 349: 2431–2441

Pham AM, Langlois RA, TenOever BR (2012) Replication in cells of hematopoietic origin is necessary for Dengue virus dissemination. PLoS Pathog 8: e1002465

Präbst K, Engelhardt H, Ringgeler S, Hübner H (2017) Basic Colorimetric Proliferation Assays: MTT, WST, and Resazurin. Methods Mol Biol 1601: 1–17

Randolph GJ, Angeli V, Swartz MA (2005) Dendritic-cell trafficking to lymph nodes through lymphatic vessels. Nat Rev Immunol 5: 617–628

Ren Z, van Andel H, de Lau W, Hartholt RB, Maurice MM, Clevers H, Kersten MJ, Spaargaren M, Pals ST (2018) Syndecan-1 promotes Wnt/β-catenin signaling in multiple myeloma by presenting Wnts and R-spondins. Blood 131: 982–994

Ribeiro CM, Sarrami-Forooshani R, Setiawan LC, Zijlstra-Willems EM, van Hamme JL, Tigchelaar W, van der Wel NN, Kootstra NA, Gringhuis SI, Geijtenbeek TB (2016) Receptor usage dictates HIV-1 restriction by human TRIM5α in dendritic cell subsets. Nature 540: 448–452

Roderiquez G, Oravecz T, Yanagishita M, Bou-Habib DC, Mostowski H, Norcross MA (1995) Mediation of human immunodeficiency virus type 1 binding by interaction of cell surface heparan sulfate proteoglycans with the V3 region of envelope gp120-gp41. J Virol 69: 2233–2239

Sarrami-Forooshani R, Mesman AW, van Teijlingen NH, Sprokholt JK, van der Vlist M, Ribeiro CM, Geijtenbeek TB (2014) Human immature Langerhans cells restrict CXCR4-using HIV-1 transmission. Retrovirology 11: 52

Sungnak W, Huang N, Bécavin C, Berg M, Queen R, Litvinukova M, Talavera-López C, Maatz H, Reichart D, Sampaziotis F et al (2020) SARS-CoV-2 entry factors are highly expressed in nasal epithelial cells together with innate immune genes. Nat Med 26: 681–687

Teng YH, Aquino RS, Park PW (2012) Molecular functions of syndecan-1 in disease. Matrix Biol 31: 3–16

Tsang NNY, So HC, Ng KY, Cowling BJ, Leung GM, Ip DKM (2021) Diagnostic performance of different sampling approaches for SARS-CoV-2 RT-PCR testing: a systematic review and meta-analysis. Lancet Infect Dis

Vanders RL, Hsu A, Gibson PG, Murphy VE, Wark PAB (2019) Nasal epithelial cells to assess in vitro immune responses to respiratory virus infection in pregnant women with asthma. Respir Res 20: 259

Wang H, Liu Q, Hu J, Zhou M, Yu MQ, Li KY, Xu D, Xiao Y, Yang JY, Lu YJ et al (2020) Nasopharyngeal Swabs Are More Sensitive Than Oropharyngeal Swabs for COVID- 19 Diagnosis and Monitoring the SARS-CoV-2 Load. Front Med (Lausanne) 7: 334

Wang N, Shi X, Jiang L, Zhang S, Wang D, Tong P, Guo D, Fu L, Cui Y, Liu X et al (2013) Structure of MERS-CoV spike receptor-binding domain complexed with human receptor DPP4. Cell Res 23: 986–993

Wang P, Nair MS, Liu L, Iketani S, Luo Y, Guo Y, Wang M, Yu J, Zhang B, Kwong PD et al (2021) Antibody resistance of SARS-CoV-2 variants B.1.351 and B.1.1.7. Nature 593: 130–135

World Health Organization, 2020a. Clinical management of COVID-19. Interim guidance 27 May 2020. World Health Organization, p. 62.

World Health Organization, 2020b. Timeline of WHO’s response to COVID-19.

Wright L, Steptoe A, Fancourt D (2020) Are we all in this together? Longitudinal assessment of cumulative adversities by socioeconomic position in the first 3 weeks of lockdown in the UK. J Epidemiol Community Health

Xia S, Liu M, Wang C, Xu W, Lan Q, Feng S, Qi F, Bao L, Du L, Liu S et al (2020) Inhibition of SARS-CoV-2 (previously 2019-nCoV) infection by a highly potent pan-coronavirus fusion inhibitor targeting its spike protein that harbors a high capacity to mediate membrane fusion. Cell Res 30: 343–355

Yuki K, Fujiogi M, Koutsogiannaki S (2020) COVID-19 pathophysiology: A review. Clin Immunol 215: 108427

Zhai Z, Li C, Chen Y, Gerotziafas G, Zhang Z, Wan J, Liu P, Elalamy I, Wang C (2020) Prevention and Treatment of Venous Thromboembolism Associated with Coronavirus Disease 2019 Infection: A Consensus Statement before Guidelines. Thromb Haemost 120: 937–948

Zhang Q, Chen CZ, Swaroop M, Xu M, Wang L, Lee J, Wang AQ, Pradhan M, Hagen N, Chen L et al (2020) Heparan sulfate assists SARS-CoV-2 in cell entry and can be targeted by approved drugs in vitro. Cell Discov 6: 80

Zhang Z, Coomans C, David G (2001) Membrane heparan sulfate proteoglycan-supported FGF2-FGFR1 signaling: evidence in support of the “cooperative end structures” model. J Biol Chem 276: 41921–41929

Zhou P, Yang XL, Wang XG, Hu B, Zhang L, Zhang W, Si HR, Zhu Y, Li B, Huang CL et al (2020) A pneumonia outbreak associated with a new coronavirus of probable bat origin. Nature 579: 270–273

Zhu N, Zhang D, Wang W, Li X, Yang B, Song J, Zhao X, Huang B, Shi W, Lu R et al (2020) A Novel Coronavirus from Patients with Pneumonia in China, 2019. N Engl J Med 382: 727–733

Zou L, Ruan F, Huang M, Liang L, Huang H, Hong Z, Yu J, Kang M, Song Y, Xia J et al (2020a) SARS-CoV-2 Viral Load in Upper Respiratory Specimens of Infected Patients. N Engl J Med 382: 1177–1179

Zou X, Chen K, Zou J, Han P, Hao J, Han Z (2020b) Single-cell RNA-seq data analysis on the receptor ACE2 expression reveals the potential risk of different human organs vulnerable to 2019-nCoV infection. Front Med 14: 185–192

